# Functional analysis of OCTN2 and ATB^0,+^ in normal human airway epithelial cells

**DOI:** 10.1101/637058

**Authors:** Bianca Maria Rotoli, Rossana Visigalli, Amelia Barilli, Francesca Ferrari, Massimiliano G. Bianchi, Maria Di Lascia, Benedetta Riccardi, Paola Puccini, Valeria Dall’Asta

## Abstract

In human, OCTN2 (SLC22A5) and ATB^0,+^ (SLC6A14) transporters mediate the uptake of L-carnitine, essential for the transport of fatty acids into mitochondria and the subsequent degradation by β-oxidation. Aim of the present study is to characterize L-carnitine transport in EpiAirway™, a 3D organotypic *in vitro* model of primary human tracheal-bronchial epithelial cells that form a fully differentiated, pseudostratified columnar epithelium at air-liquid interface (ALI) condition. In parallel, Calu-3 monolayers grown at ALI were used as comparison. In EpiAirway™, ATB^0,+^ was highly expressed and functional on the apical side while OCTN2 transporter was active on the basolateral side. Calu-3 cells showed a different pattern of expression and activity for ATB^0,+^: indeed, L-carnitine uptake on apical side was evident in Calu-3 at 8 days of culture but not in fully differentiated 21d ALI culture. As both ATB^0,+^ and OCTN2, beyond transporting L-carnitine, have a significant potential as delivery systems for drugs, the identification of these transporters in EpiAirway™ can open new fields of investigation in the studies of drug inhalation and pulmonary delivery.

## INTRODUCTION

L-Carnitine (β-hydroxy-γ-trimethylaminobutyrate) is a small highly polar zwitterionic molecule essential in the transfer of activated long-chain fatty acids across the inner mitochondrial membrane in a series of reactions called the “carnitine shuttle”, so that they can undergo degradation by β-oxidation [1]. Besides its key role in energy metabolism, several studies provide evidence that L-carnitine also functions as a cytoprotector by promoting cell resistance and antiapoptotic pathways, as well as by enhancing antioxidative resources [2,3,4]. In humans, the transport of L-carnitine across the cell membrane is mediated by OCTN1 (SLC22A4), OCTN2 (SLC22A5) and ATB^0,+^ (SLC6A14).

OCTN2 operates a Na^+^-dependent, high-affinity (Km is in the range of 10-20 μM) transport of L-carnitine, the physiological substrate, and other carnitine derivatives, as well as a Na^+^-independent transport of organic cations [5,6]. In polarized epithelia such as intestine and kidney OCTN2 is located in the apical membrane of the cells [7] where it is involved in intestinal absorption and renal reabsorption. The transporter is also expressed in human macrophages, where it has been identified as a novel target gene of the mTOR-STAT3 axis [8]. Moreover, OCTN2 expression has been demonstrated in liver, heart, testis, skeletal muscle, lung and brain, sustaining a role for the transporter in the systemic distribution of carnitine [9]. The role of OCTN2 in the modulation of carnitine availability is clearly evidenced by the autosomal recessive disorder “Systemic primary carnitine deficiency” (SPCD; OMIM 212140) in which OCTN2 mutations strongly compromise the function of several tissues [1] and lead to severe clinical signs, such as respiratory insufficiency, vomiting, progressive cardiomyopathy, skeletal myopathy, hypoglycemia, and hyperammonemia. Due to the genetic defect, dietary L-carnitine is poorly absorbed in the intestinal tract and lost in the urines because of the defective renal reabsorption from the glomerular filtrate; this reduces circulating carnitine levels and decreases intracellular carnitine accumulation with consequent impairment of fatty acid oxidation.

Contrary to OCTN2, the physiological role of OCTN1 is thus far not unequivocally defined. The transporter, specific for ergothioneine [10], is endowed with a only low affinity for L-carnitine. However, the group of Longo N. demonstrated some years ago that the expression of human OCTN1 in transfected CHO (Chinese Hamster Ovary) cells fails to cause any significant increase in L-carnitine transport [11]; similarly, we recently demonstrated that, despite its expression in human bronchial A549 and BEAS-2B cells, OCTN1 in not involved in carnitine transport in these models [8].

The other transporter involved in carnitine absorption is ATB^0,+^, a system responsible for the Na^+^/Cl^-^-dependent influx of neutral and cationic amino acids. This transporter has a low affinity (Km = 800 μM) for L-carnitine, but an high concentrative capacity, being energized by the transmembrane gradients of Na^+^ and Cl^-^, as well as by membrane potential [12]. Consistent with functional studies, ATB^0,+^ is expressed in the lung and intestine under normal conditions [13,14] where it is supposed to be mainly involved in nutrient uptake, due to its broad specificity and concentrative transport mechanisms [15,16].

In our previous study we performed a characterization of L-carnitine transport in different human airway epithelial models (A549, Calu-3, NCl-H441, and BEAS-2B), demonstrating that OCTN2 is the only L-carnitine transporter in A549 and BEAS-2B cells, while both OCTN2 and ATB^0,+^ are operative in Calu-3 and NCl-H441 [17]. It is noteworthy, however, that these models are immortalized or transformed cell lines and criticisms exist as their biological functions can differ from those of primary differentiated human airway epithelial cells; moreover, the biological features of these cells may be modified under culture conditions and, in particular for Calu-3, evidences established a clear dependence on culture time and passage number for optimal barrier integrity, mucous secretion, and P-gp expression [18].

The purpose of the present study is, hence, to characterize L-carnitine transport in an in vitro model of normal human respiratory cells; to this end, EpiAirway™ a 3D organotypic model of human mucociliary airway epithelium consisting of normal, human-derived differentiated tracheal/bronchial epithelial cells, has been employed and compared with Calu-3 cells grown under air-liquid interface (ALI) condition.

## METHODS

### EpiAirway™ primary cultures

EpiAirway™ tissues (AIR-200-PE6.5), supplied by MatTek Corporation (Ashland, MA), were used. Cultured on microporous membrane inserts at the air-liquid interface (ALI), EpiAirway™ tissues recapitulates aspects of the *in vivo* microenvironment of the lung [19,20]; this system is, indeed, produced from primary human tracheal-bronchial epithelial cells that form a fully differentiated, pseudostratified columnar epithelium containing mucus-producing goblet cells, ciliated cells and basal cells. Upon arrival, tissue inserts were transferred to 24-well plates containing 600 μl of the AIR 200-M125 medium and equilibrated overnight at 37°C and 5% CO_2_; medium at the basolateral side was, then, renewed every day, while apical washes for mucus removal were performed employing the solution provided by the manufacturer. Cultures from four different healthy donors were employed.

### Calu-3 cell line

Calu-3 cells (American Type Culture Collection), obtained from a human lung adenocarcinoma and derived from serous cells of proximal bronchial airways, were cultured in Eagle’s Minimum Essential Medium (EMEM) supplemented 10% fetal bovine serum (FBS), sodium pyruvate (1 mM) and 1% penicillin/streptomycin. Cells between passages 25-30 were routinely cultured under physiological conditions (37.5 °C, 5% CO_2_, 95% humidity) in 10-cm diameter dishes. For the experiments, Calu-3 cell monolayers were grown at air-liquid interface (ALI). To this end, cells were seeded onto Transwell polyester inserts (0.33 cm^2^, 0.4 μm pore size; Falcon) at the density of 10^5^ cells/insert; the apical medium was removed 24 hours after seeding, while basolateral medium was renewed every other day. The monolayers were allowed to differentiate under ALI condition over 8 or 21 days.

### Carnitine uptake

Carnitine transport was measured both at apical and at the basolateral side. After two washes in pre-warmed transport buffer (Earle’s Balanced Salt Solution (EBSS) containing (in mM) 117 NaCl, 1. 8 CaCl_2_, 5.3 KCl, 0.9 NaH_2_PO_4_, 0.8 MgSO_4_, 5.5 glucose, 26 Tris/HCl, adjusted to pH 7.4), cells were incubated in fresh transport buffer (100 μl apical and 600 μl basolateral) containing L-[^3^H]carnitine (2 μCi/ml). For a sodium-free EBSS, 117 mM NaCl was replaced with equimolar N-methyl-D-glucamine chloride. Where indicated, the inhibitors were added to the transport buffer at the indicated concentrations. After 30 min, transport buffer was removed, and the experiment terminated by two rapid washes (< 10 s) in ice-cold 300 mM urea. The filter was, then, detached from the insert and the ethanol soluble pool was extracted from monolayers; radioactivity in cell extracts was determined with MicroBeta2 liquid scintillation spectrometer (Perkin Elmer, Italy). Protein content in monolayers was determined directly in the filters using a modified Lowry procedure [21]. L-carnitine uptake is expressed as pmol/mg of protein, as indicated. In order to discriminate the contribution of OCTN2 and ATB^0,+^ transporters, carnitine transport was measured either in the absence or in the presence of betaine or arginine employed as inhibitors of OCTN2 and ATB^0,+^, respectively [17]. Hence, the activity of ATB^0,+^ is calculated from the difference between total transport and that measured in the presence of arginine, while the activity of OCTN2 results from the difference between total transport and that measured in the presence of betaine.

### RT-qPCR analysis

Total RNA was isolated with GeneJET RNA Purification Kit and reverse transcribed with RevertAid First Strand cDNA Synthesis Kit (Thermo Fisher Scientific, Italy). The expression of SLC6A14/ATB^0,+^ was measured using specific forward/reverse primers (5’GCTGCTTGGTTTTGTTTCTCCTTGGTC3’ and 5’GCAATTAAAATGCCCCATCCAGCAC3’) and SYBR™ Green PCR Master Mix (Thermo Fisher Scientific); the amount of SLC22A5/OCTN2 and that of the housekeeping gene RPL15 (Ribosomal Protein Like 15) were, instead, monitored employing specific TaqMan^®^ Gene Expression Assays (Thermo Fisher Scientific; Cat# Hs00929869_m1 and Hs03855120_g1, respectively), according to the manufacturer’s instructions. The amount of the genes of interest is expressed relatively to that of the housekeeping gene RPL15, using the formula 1000 × 2^ΔCt^ (ΔCt = Ct_RPL15_-Ct_gene of interest_).

### Transepithelial electrical resistance of cell layers

Before each experiment, the integrity of the layers was assessed by measuring the transepithelial electrical resistance (TEER) using an epithelial voltohmmeter with chopstick electrodes (World Precision Instruments, Stevenage, UK). TEER values of ~ 180 Ω · cm^2^ and ~ 160 Ω · cm^2^ were measured for EpiAirway^TĨVI^ and Calu-3 cells. The flux of ^14^C-mannitol yielded permeability values of about 4.5 · 10^-7^ cm/sec and 2.1· 10^-7^ cm/sec, respectively.

### Immunofluorescence staining and analysis with confocal laser scanning microscopy (CLSM)

Calu-3 monolayers and EpiAirway™ tissues were rinsed in phosphate buffered saline (PBS), then fixed with ice cold methanol for 7 min or with 3.7% paraformaldehyde at RT for 15 min, respectively. Cells were then permeabilized with a 20 min incubation in 0.2% Triton X-100 in PBS. After 1h in blocking solution (5% of bovine serum albumin in PBS) at 37 °C, cells were incubated overnight at 4 °C with the polyclonal antibody ATB^0,+^ (1:100, Sigma-Aldrich), then washed with PBS and further incubated for 45 min with Alexa Fluor 488 antibody (1:400, Abcam). After three rinses, cells were incubated with propidium iodide solution (Cell Signaling, EuroClone) for 10 min at 37 °C, so as to stain nuclei; control filters were stained with propidium iodide in the absence of primary antibodies, so as to obtain only nuclei signal. After staining procedures, the filters were detached from the culture inserts and mounted on microscope slide glass slides with fluorescence mounting medium (FluorSave Reagent, Calbiochem). Samples were observed with a Zeiss^®^ 510 LSM Meta confocal microscope through a 63x (NA 1.4) oil objective, adopting a multi-track detection mode. Excitation at 488 nm and emission recorded through a 505-530 nm band pass barrier were used for the detection of the transporters; excitation at 543 nm and emission recorded through a 580-630 nm band pass barrier filter were adopted for cell nuclei. The two signals were rendered with scales of green and red for transporters and nuclei, respectively. Vertical sections were obtained with the function Display – Cut (Expert Mode) of the LSM 510 confocal microscope software (Microscopy Systems, Hartford, CT). Reconstructions were performed from z-stacks of digital images, processed with the Axiovision module inside 4D release 4.5 (Carl Zeiss), applying the shadow or the transparency algorithm.

### Statistical analysis

GraphPad Prism 7 (GraphPad Software, San Diego, CA, US) was used for statistical analysis. P values were calculated with a two tailed Student’s t-test for unpaired data; p < 0.05 was considered significant.

### Materials

Fetal bovine serum was purchased from EuroClone (Milano, Italy). Carnitine-L-[N-methyl-^3^H]HCl (80 Ci/mmol) was obtained from Perkin-Elmer (Milano, Italy). Sigma-Aldrich (Milano, Italy) was the source of the antibodies against ATB^0,+^ and, unless otherwise specified, of all other chemicals.

## RESULTS

In order to address the specific transporters involved in L-carnitine uptake in EpiAirway™ cells, 30-min uptake of 1 μM carnitine was performed at both apical and basolateral side in the absence and in the presence of sodium and in the presence of specific transporters inhibitors (Figure 1). Carnitine influx was totally sodium-dependent at both culture sides, being the transport almost completely suppressed in the absence of the cation. On the apical compartment, L-carnitine uptake was fully inhibited by arginine, which share with carnitine ATB^0,+^ transporter, while betaine, substrate of OCTN2, was completely ineffective. Otherwise, at basolateral side, while betaine markedly inhibited carnitine transport, arginine was completely ineffective. Overall, these findings indicate that L-carnitine transport in EpiAirway™ cell system is mediated by ATB^0,+^ transporter on the apical side and by OCTN2 in the basolateral side.

**Figure 1.**
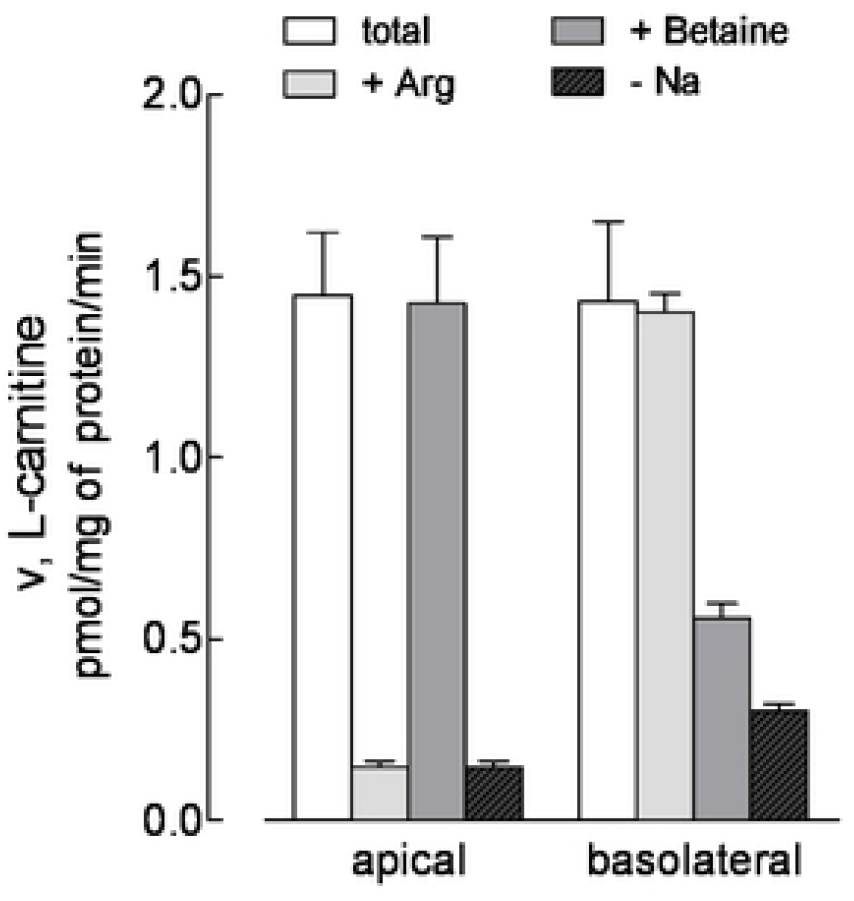
Monolayers of EpiAirway™ cells grown under air-liquid interface (AU) conditions were washed with EBSS (Na^+^-present, total uptake) or Na^+^-free EBSS (-Na^+^), as indicated. L-Carnitine uptake was then assayed with 30-min incubation in the same buffer containing [^3^H]carnitine (1 μM; 2 μCi/ml), either added to the apical or to the basolateral compartment. Where indicated, arginine (2 mM) or betaine (2 mM) were present during the transport assay. The intracellular L-carnitine was determined as described in section Material and Methods. Bars represent the mean ± S.D. of three independent determinations in a representative experiment that, repeated twice, gave comparable results.

The same functional analysis was then applied to Calu-3 cells cultured under Air-liquid Interface (ALI) conditions for 21 d, so as to obtain a comparison between the two cell models (Figure 2, panel A). Unexpectedly, carnitine uptake at the apical side was very low, not significantly inhibited by the presence of either arginine or betaine, and not affected by the absence of sodium. At the basolateral side, on the contrary, transport data were comparable with those of EpiAirway™ cells, being carnitine influx totally sodium-dependent and completely inhibited by betaine and not by arginine. In order to better address the differences observed between the two models, the same analysis was repeated in Calu-3 cells cultured under ALI conditions for shorter time (8 d). As shown in Figure 2, panel B, L-carnitine transport measured at the apical side was higher than in Calu-3 cultured for 21 d and comparable to that of EpiAirway™, with the same inhibition pattern; similarly, also data obtained at the basolateral side overlapped with those of primary normal cells. Transport data were consistent with the expression of the different transporters at gene level: the mRNA coding for SLC22A5/OCTN2 was, indeed, equally expressed in EpiAirway™ and in Calu-3, both cultured for 8 and 21 d; SLC6A14/ATB^0,+^, on the contrary, was maximally detectable in EpiAirway™, expressed to a lesser extent in Calu-3 at 8 d and much less abundant after 21 d of culture (Figure 3).

**Figure 2.**
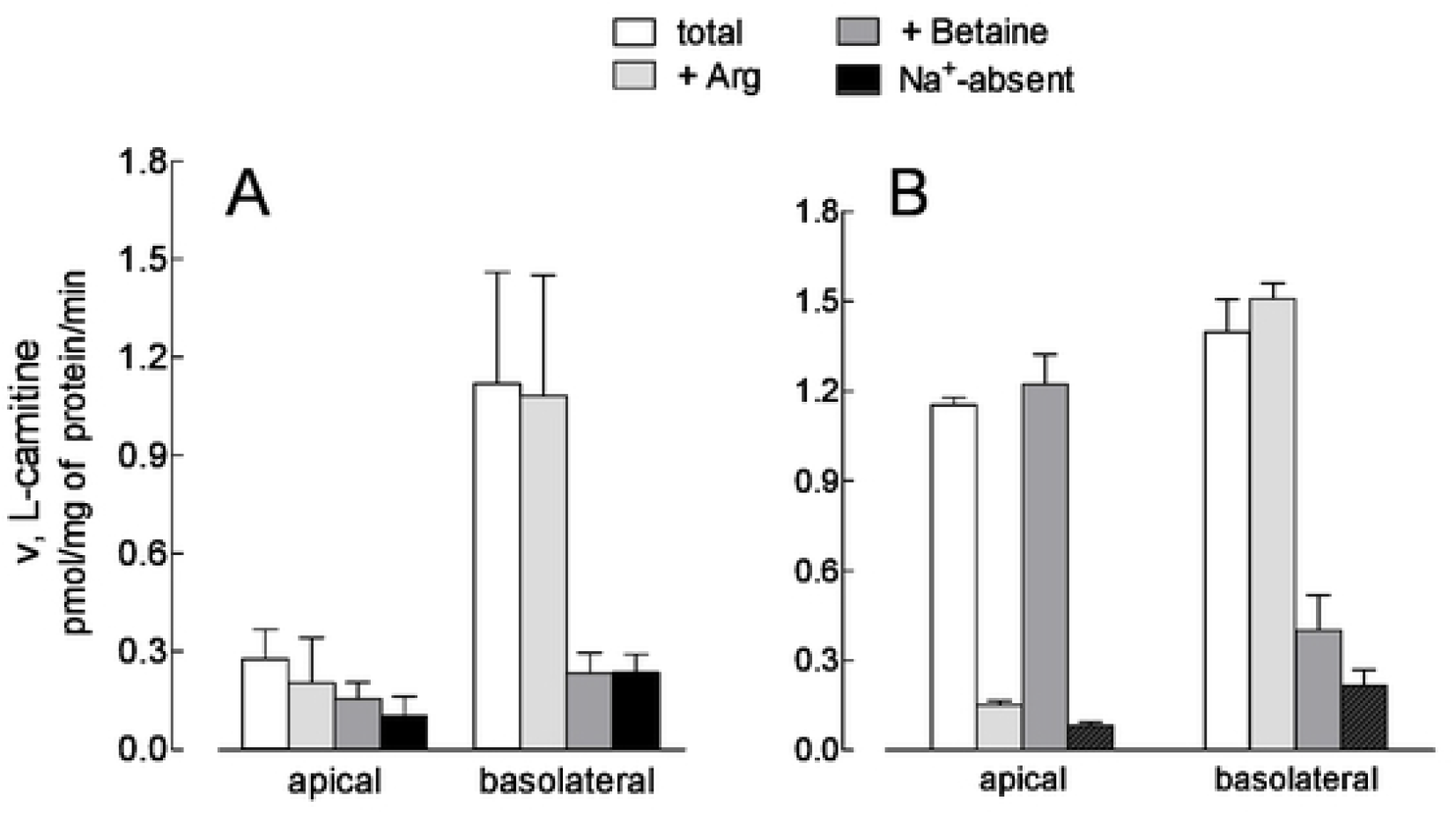
Monolayers of Calu-3 cells grown under air-liquid interface (AU) conditions for 21 d (panel A) or 8 d (panel B) were washed with EBSS (Na^+^-present, total uptake) or Na^+^-free EBSS (-Na^+^), as indicated. L-Carnitine uptake was assayed with 30-min incubation in the same buffer containing [^3^H]carnitine (1 μM; 2 μCi/ml), either added at the apical or at the basolateral side. Where indicated, arginine (2 mM) or betaine (2 mM) were present during the transport assay. Bars represent the mean ± S.D. of three independent determinations in a representative experiment, repeated three times with comparable results.

**Figure 3.**
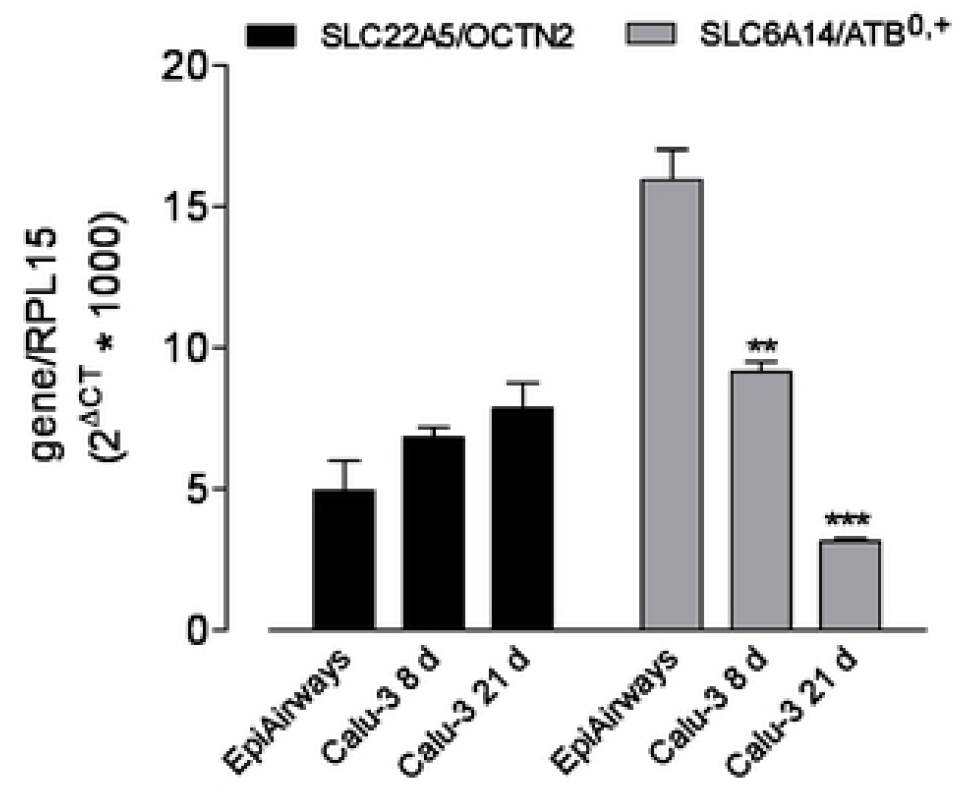
mRNA levels for SLC22A5/OCTN2 and SLC6A14/ ATB^0,+^ were determined in EpiAirway™ and in Calu-3 cultured under ALI conditions for 8 d or 21 d (as indicated) by means of quantitative RT-PCR analysis. The expression of the gene of interest was shown after normalization for that of the housekeeping gene (RPL15). Data are means ± SEM of three experiments, each performed in duplicate, **p < 0.01, ***p < 0.001 vs EpiAirway™.

The expression of ATB^0,+^ was lastly assessed at protein level by means of immunocytochemistry. This approach confirmed a clear-cut distribution of the transporter on plasma membranes at the apical side of both EpiAirway™ (Figure 4) and Calu-3 monolayers cultured for 8 d (Figure 5, right panel); on the contrary, the protein was only barely detectable in Calu-3 after 21 d of culture (left panel).

**Figure 4.**
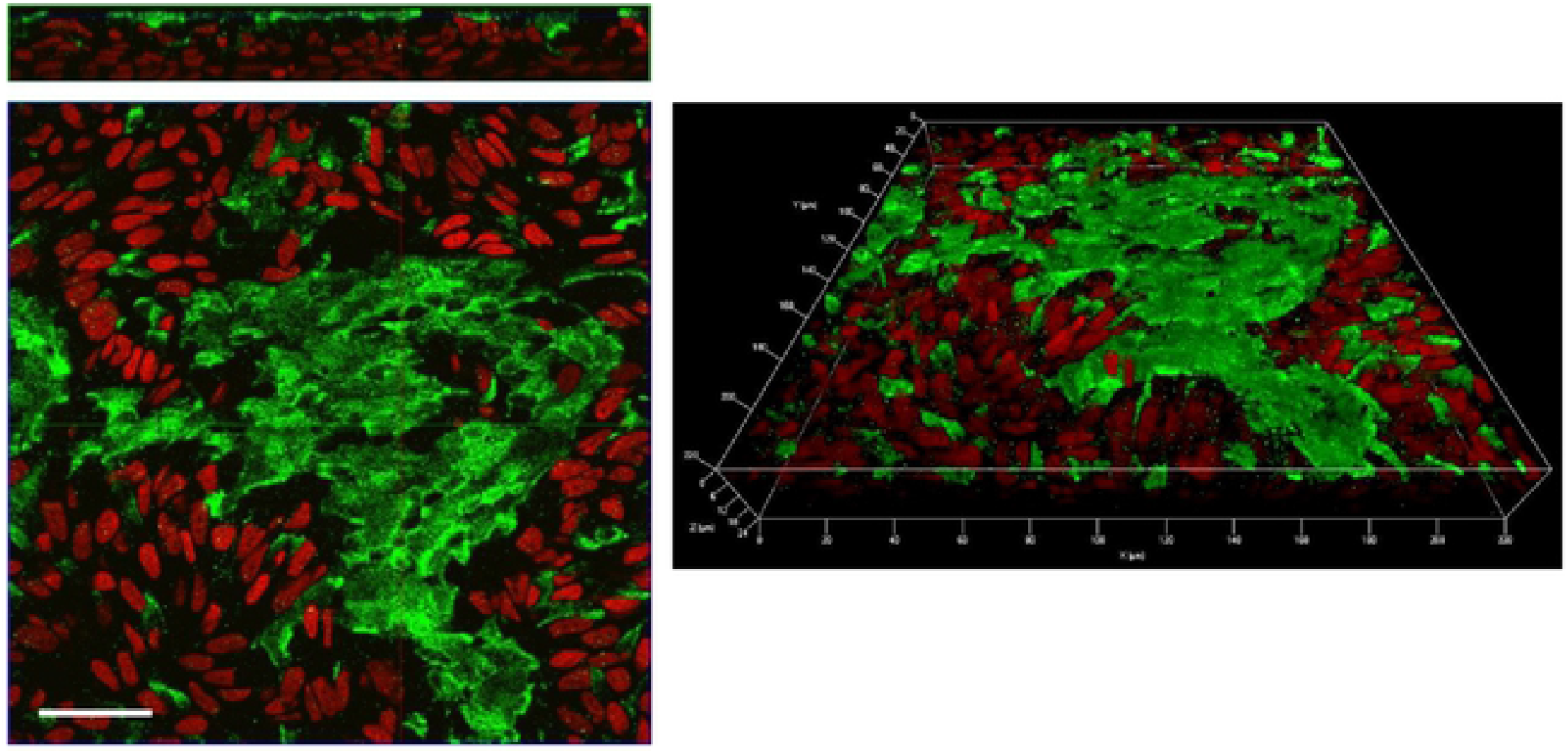
Confocal laser scanning microscopy of EpiAirway™ layers immunolabelled for ATB^0,+^ transporter (green). Cells were counterstained with propidium iodide (red). Left: single plane acquired in correspondence of the apical membrane. At the top, a vertical section of the plane is shown. Right: 3D reconstruction of z-stacks confocal images; about 30 horizontal sections were acquired. Bar, 50 μm

**Figure 5.**
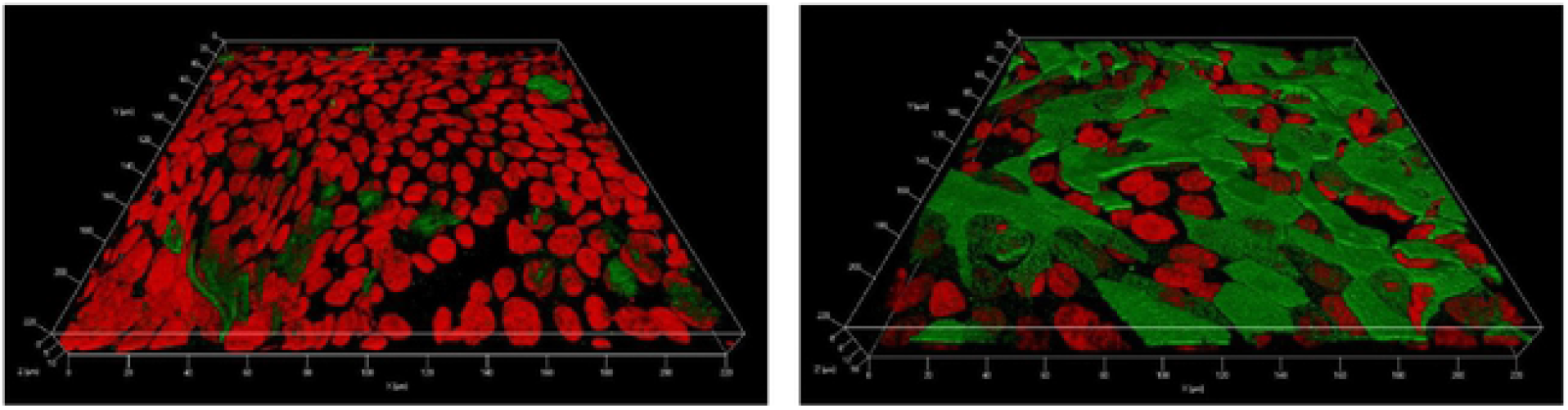
3D reconstructions of z-stacks of confocal images of Calu-3 monolayers immunolabelled for ATB^0,+^ transporter (green). Cells were counterstained with propidium iodide (red). Cells were cultured under ALI condition for 21 days (left) or 8 days (right). The images were obtained through the acquisition of about 20 horizontal sections.

## DISCUSSION

This study is the first to evaluate OCTN2 and ATB^0,+^ transporters in an organotypic model of human airway epithelium like EpiAirway™. To date, most of the studies dealing with the expression of transporters have used cell lines such as Calu-3, BEAS-2B and 16HBE14- cells as models of human bronchial epithelium. The main criticism lies in the fact that, being lines, cells are immortalized or transformed, and can therefore show biological functions that deviate from those of primary respiratory cells obtained *ex vivo* from tissues [22]. In recent years, 3D systems of human primary airway cells have been developed, such as the EpiAirway™, which appear promising and innovative since, grown at ALI, they closely resembles the epithelium *in vivo.* Indeed, EpiAirway™ tissues consist of a pseudostratified epithelium that contain the differentiated cell types found in the respiratory epithelium like mucus-producing goblet cells, ciliated and basal cells.

However, to date, little information is available on the biophysical characteristics of these cells; in particular minimal information is available evaluating the identification of membrane transporters responsible of fluxes of nutrients, such as amino acids, sugar, vitamins, and other substances like drugs.

In this study the functional activity of ATB^0,+^ and OCTN2 transporters in EpiAirway ™ was compared with that of Calu-3 monolayers, one of the reference model in the studies of transporter proteins and drug absorption [23]. In EpiAirway™, the functional analysis of the transport of L-carnitine, substrate of both ATB^0,+^ and OCTN2, revealed that ATB^0,+^ is present at apical level while OCTN2 at the basolateral side. As far as ATB^0,+^ is concerned, data of transport analysis are fully in agreement with the immunocytochemical staining which showed a strong positivity of the protein at plasma membrane at the apical side. As far as the biologiocal role of this transporteer, it has been suggested that ATB^0,+^ is expressed at places where the body interfaces with microbes, such as lung and colon and it may be involved in reducing available nutrients to bacteria [14]. Accordingly, the transporter is upregulated in inflammatory states, such as ulcerative colitis, Crohn’s disease and colon cancer [14]. Moreover, genome-wide association studies has been recently identified ATB^0,+^ (SLC6A14) as a genetic modifier of lung disease severity in cystic fibrosis, providing a mechanism by which it regulates Pseudomonas aeruginosa attachment to human bronchial epithelial cells [24]. Compared to Calu-3 monolayers, a close similarity between the two cell models is observed only when Calu-3 layers are maintained at ALI for 8 days. Indeed, the uptake of carnitine through ATB^0,+^ as well as its expression, are no longer appreciable when the cultures of Calu-3 cells are prolonged to 21 days, when the monolayers are fully differentiated. This finding, hence, confirms that the biological features of these cells may be modified by the culture conditions, and includes the expression of ATB^0,+^ to the list of parameters that vary along with the duration of the culture. As far as OCTN2 expression, here we show that carnitine transport through OCTN2 is similar in EpiAirway™ and Calu-3 where it appears localized on the basolateral membranes of both cell models. Our functional data are not in accordance with an immunohistochemistry analysis recently performed in lung tissues from healthy and COPD patients [25]: in this study, indeed, OCTN2 is founded on the apical and lateral side of the epithelial cells in the bronchial region and in cells lining the bronchioles. Since our findings do not highlight any transport activity through OCTN2 at the apical side, further investigations are necessary to ascertain the discrepancy between the positivity of immunohystochemical staining and the actual functional activity.

Beyond transporting L-carnitine, ATB^0,+^ and OCTN2 have significant potential as delivery systems for amino acid-based drugs and prodrugs [13,26]; OCTN2 in particular interacts with inhaled drugs e.g. muscarinic antagonists and β-adrenergic agonists cationic bronchodilators [27]; moreover, this transporter offer an efficient means to deliver drug or drug-loaded nanoparticles conjugated to carnitine [28]. Overall, the identification of the ATB^0,+^ and OCTN2 transporters in the EpiAirway™ can open new fields of investigation in the studies of drug inhalation and pulmonary delivery.

## CONCLUSIONS

In conclusion, in EpiAirway™ cells carnitine is transported through ATB^0,+^ transporter at apical side and OCTN2 transporter at the basolateral side. A similar pattern of transporters is present only in Calu-3 monolayers cultured at ALI condition for short time; indeed when the culture was prolonged for 21 d, the activity of ATB^0,+^ was no more detectable.

## Abbreviations

ALI: air-liquid interface
ATB^0,+^: amino acid transporter B^0,+^
3D: three-dimensional
EBSS: Earle’s Balanced Salt Solution
OCTN: novel organic cation transporter
PBS: phosphate-buffered saline solution
TEER: transepithelial electrical resistance.

